# Molecular anatomy and plasticity of the long noncoding RNA HOTAIR

**DOI:** 10.1101/444331

**Authors:** Rachel Spokoini-Stern, Dimitar Stamov, Hadass Jessel, Lior Aharoni, Heiko Haschke, Jonathan Giron, Ron Unger, Eran Segal, Almogit Abu-Horowitz, Ido Bachelet

**Author notes:** These authors contributed equally to this work (Rachel Spokoini-Stern, Dimitar Stamov). Address for correspondence: 8 Hamada St. Rehovot, 7670308, Israel.

## Abstract

Long noncoding RNA molecules (lncRNAs) are estimated to account for the majority of eukaryotic genomic transcripts, and have been associated with multiple diseases in humans. However, our understanding of their structure-function relationships is scarce, with structural evidence coming mostly from indirect biochemical approaches or computational predictions. Here we describe the hypothetical molecular anatomy of the lncRNA HOTAIR (HOx Transcript AntIsense RNA) inferred from direct, high-resolution visualization by atomic force microscopy (AFM) in nucleus-like conditions at 37 degrees. Our observations reveal that HOTAIR has a distinct anatomy with a high degree of plasticity. Fast AFM scanning enabled the quantification of this plasticity, and provided visual evidence of physical interactions with genomic DNA segments. Our report provides the first biologically-plausible hypothetical description of the anatomy and intrinsic properties of HOTAIR, and presents a framework for studying the structural biology of lncRNAs.

---

Long Non Coding RNA (lncRNA) molecules are defined as RNA transcripts longer than 200 nucleotides, that lack an evident ORF and are transcribed by RNA polymerase II. These large transcripts are often also polyadenylated and spliced. Thousands of lncRNAs are annotated to date, but only a few have been studied and defined as functional. Functional lncRNAs are involved in almost every stage of gene expression^1–4^ and have been implicated in a variety of diseases such as cancer and neurodegenerative disorders.

Despite their abundance and emerging importance, our knowledge concerning lncRNA structure is poor. Existing information on the structure of large RNA molecules in general is scarce, with less than 7% of all RNA structures in the Protein Data Bank being in the size range between 200 and 5,000 nucleotides, most of these being ribosomal RNA subunits^5^ studied by X-ray crystallography. The structure of MALAT1 has been resolved by crystallography^6^, but available information on other lncRNA structures derives mostly from indirect methods. For example, the structure of SRA^7^ and HOTAIR^8^ were depicted through biochemical methods. Another direction taken to further study lncRNAs such as XIST^9^ and GAS5^10^ was domain analysis. On the other hand, it has recently been suggested, using a statistical approach, that lncRNAs may not have a structure at all^11^. This discrepancy may result from lack of solid structural information, and its resolution could shed light on the biology of this important class of molecules.

In this work we aimed to obtain such information using one of the most studied lncRNAs, HOTAIR, as a test case. HOTAIR has been shown to bind PRC2 and LSD1^12^ to drive chromatin modification at specific genomic sites^13^, thus playing a key role in genome silencing. HOTAIR DNA binding sites are focal, specific and numerous, implying HOTAIR as a silencing selector element pinpointing genomic locations to modification^13^. HOTAIR was shown to be required for the epithelial-to-mesenchymal transition^14,15^, thereby defined as an oncogene and a negative prognostic marker in various cancers^16–18^.

Our central tool in this study was atomic force microscopy (AFM). AFM has been used to study nucleic acid structures in both fluid and air^19^, including the genomic RNA of human immunodeficiency virus (HIV)-1^20^. While AFM is limited in resolution compared with X-ray crystallography or NMR, it enables direct visualization by physically probing native, large molecules under biological conditions. AFM also allows statistical analysis of structurally-diverse molecules, as previously suggested to be the case for lncRNAs^21^.

We first generated HOTAIR molecules by in-vitro transcription (IVT), using multiple templates and multiple IVT systems in order to avoid method-biased observations. The resulting transcripts were analyzed by gel electrophoresis and RNA-seq and found to be intact and to fully map to the HOTAIR gene sequence in a human reference genome **(Supplementary notes 1 & 2)**. HOTAIR molecules were then scanned by AFM in fluid, under conditions that mimic the chemical environment of the nucleus as reliably as possible^22,23,24^ and at 37 °C **(Supplementary note 3)**.

AFM scanning demonstrated that under these conditions, HOTAIR molecules assume a distinct anatomy **(Fig. 1A,B)**, based on a 4-limbed body which ends in a branched U-shaped motif, which we termed the U-module. Visualization of HOTAIR by Cryo-EM showed the same anatomy and flexibility observed in AFM **(Fig. 1C, Supplementary note 4)**. This anatomy was reproduced in 7 independent synthesis and scanning repeats. In contrast, a random RNA transcript formed by scrambling HOTAIR sequence formed indistinct shapes and aggregates, with some even not folding **(Supplementary note 5)**. The archetypal HOTAIR anatomy could be reliably assigned to ~2/3 of the observed objects that were intact based on size, with excluded ones being either clear but eccentric or too vague to be reliably assigned. The presence of eccentric forms could reflect that not all HOTAIR products fold properly in the nucleus and are subsequently dysfunctional, however the folding of HOTAIR in the nucleus could be facilitated by yet unidentified cellular factors. In addition to the whole molecule, the functional HOTAIR modules suggested in previous studies^12,13,25,26^ were also scanned, showing distinct structures **(Supplementary note 6, Movie S1-S3)**.

**Figure 1.**
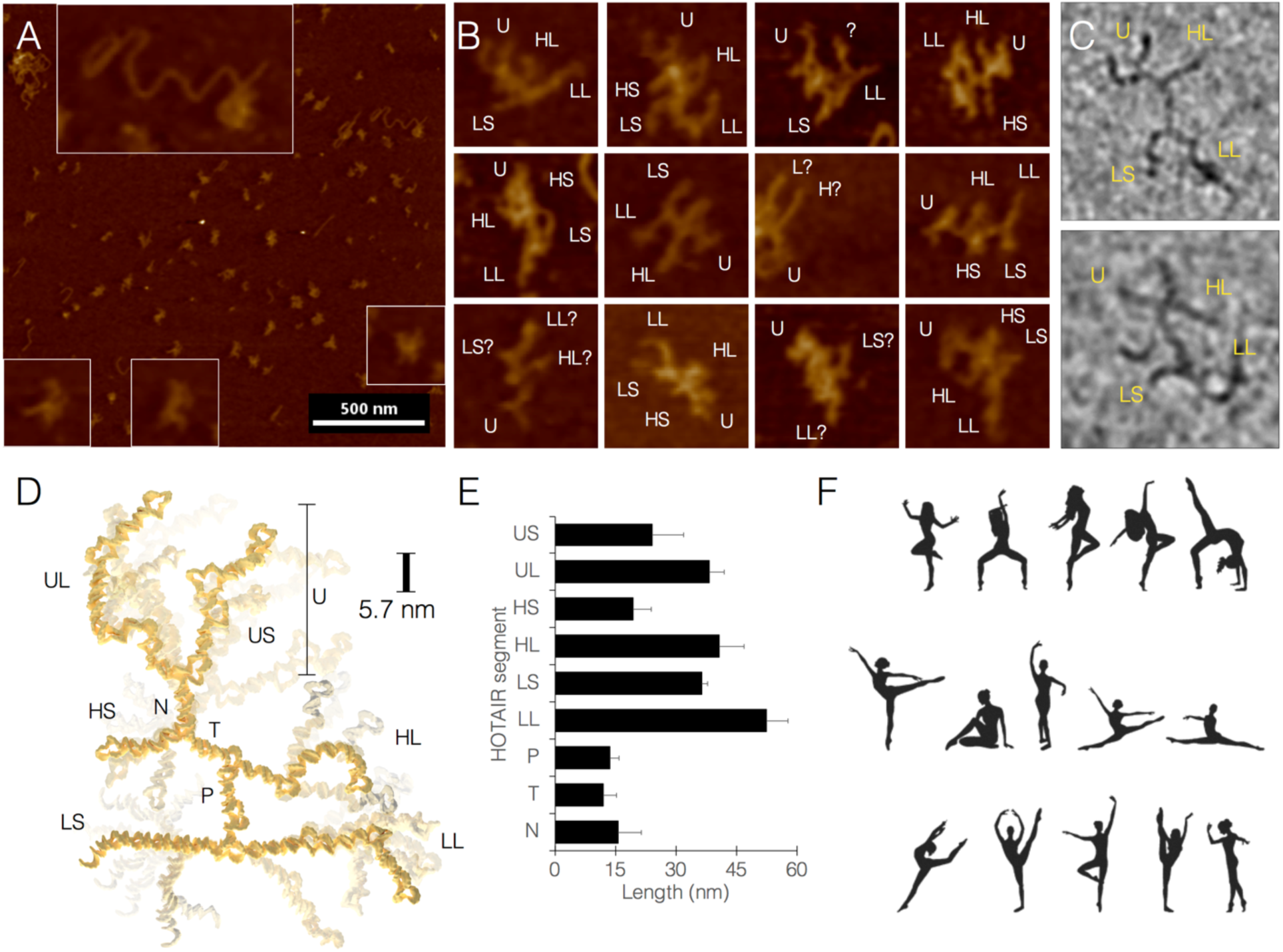
Visualization and modeling of the molecular anatomy and plasticity of HOTAIR by AFM. A, field imaged at lower magnification showing free HOTAIR molecules (bar = 500 nm). Representative objects that can be reliably assigned are enlarged in square frames. Rectangular frame shows HOTAIR molecule interacting with DNA. B, higher magnification images of HOTAIR molecules showing their distinct anatomy. Frame height/width = 70 nm. C, representative cryo-EM images of HOTAIR. Frame height/width = 70 nm. D, proposed molecular model of HOTAIR, divided into 9 structural segments (UL, U long; US, U short; N, neck; HS, hand short; HL, hand long; T, torso; P, pelvis; LS, leg short; LL, leg long). E, HOTAIR plasticity expressed as variation in segment lengths. F, the human figures frozen in various configurations are analogous to the concept of HOTAIR as a molecule with a distinct anatomy but also high flexibility.

Our HOTAIR model divides the molecule into 9 structural segments, which we name as follows: neck (N), torso (T), pelvis (P), leg long (LL), leg short (LS), hand long (HL), hand short (HS), U-module long (UL), U-module short (US) **(Fig. 1D)**. Total length of all segments across multiple samples was constrained to 252±9.4 nm, but the observed variance within segments was 5% to 50% of the mean segment length (shorter segments exhibited higher variance) **(Fig. 1E)**. The high degree of flexibility in our proposed model of HOTAIR may seem contradictory to the conventional meaning of a molecule having a structure, however this notion is contained within a set of objects that are well-defined anatomically but are globally flexible. A useful analogy is shown here by human dancers frozen in various configurations **(Fig. 1F)**, who also combine these two properties.

The molecular dimensions of HOTAIR were measured by an algorithm that combined the absolute size calculated by the AFM with a dsDNA molecule as an internal size reference. We chose to use a dsDNA molecule termed HOTAIR-binding DNA 1 (HBD1)^18^, which we synthesized for this study **(Supplementary note 7)**. HBD1 is a 433 bp molecule existing preferentially at the B-DNA geometry (helical rise of 3.5 Å per base with 10.5 bp/turn, yielding a longitudinal density of 3 bp/nm) under the study conditions, thus mapping to 144 nm in length. In contrast, HOTAIR, a 2,158^27^ nt RNA molecule which is mostly dsRNA, preferentially exists at the A-DNA geometry (helical rise of 2.6 Å per base with 11 bp/turn, yielding a longitudinal density of 4.23 bp/nm). Our measurements **(Supplementary note 8)** showed a mean length of 142 nm for HBD1, and 252 nm for HOTAIR, the latter translating to 1,066 base pairs of dsRNA, which theoretically unfold to 2,132 nt of ssRNA. Taken together, this calculation represents a ~1.5% error in molecular measurements by AFM under the study conditions.

It is critical to note that our aim in this study was to report *intrinsic* properties of HOTAIR, particularly its ability to interact with genomic DNA, dissociated from the suggested role of auxiliary proteins in this function, which is still unclear. For example, a recent study reported that genome targeting by HOTAIR is independent of at least one specific protein it interacts with, EZH2, although it did not rule out other proteins such as those that are part of the complex LSD1^13^. With that in mind, our aim here was first to describe the structural ‘baseline’ of HOTAIR, on top of which future investigations of the functional complexes it forms inside the cell could be carried out.

To this end, two dsDNA sequences from a previous study^13^ were used, one that was found to highly associate with HOTAIR and one that was found not to, termed HOTAIR-binding DNA 1 (HBD1) and HBD4, respectively **(Supplementary note 7)**. HBD1 and HBD4 were allowed to interact with HOTAIR for short times, up to 30 min at various HOTAIR:HBD ratios, and fast AFM scanning was used to count temporally-and positionally-defined interactions (remaining bound at the same position along several scan frames). AFM scans demonstrated clear physical interaction between HOTAIR and DNA **(Fig. 2A-C, Movies S4-S6)**. Several interesting features of these interactions were observed. First, assignments based on plausible configurations combined with length measurements revealed that HOTAIR:DNA interactions appear to be mediated by the U-module **(Fig. 2D-E)**. In some cases a second interaction is seen mediated by an H segment **(Fig. 2A,C)**. Second, the 1200 nt domain of HOTAIR, previously shown to associate with DNA^26^, exhibited this ability under the study conditions, suggesting that the U-module maps to this specific domain **(Fig. 2F, Movie S7)**. Third, the occupancy of HBD1 by HOTAIR was ratio-dependent **(Fig. 2G)**, suggesting a real biological phenomenon. Finally, the occupancy of HBD1 by HOTAIR was 3.2-fold higher than that of HBD4 (46 occupied out of 60 total HBD1 molecules counted vs. 16 occupied out of 66 total HBD4 counted) **(Fig. 2H)**.

**Figure 2.**
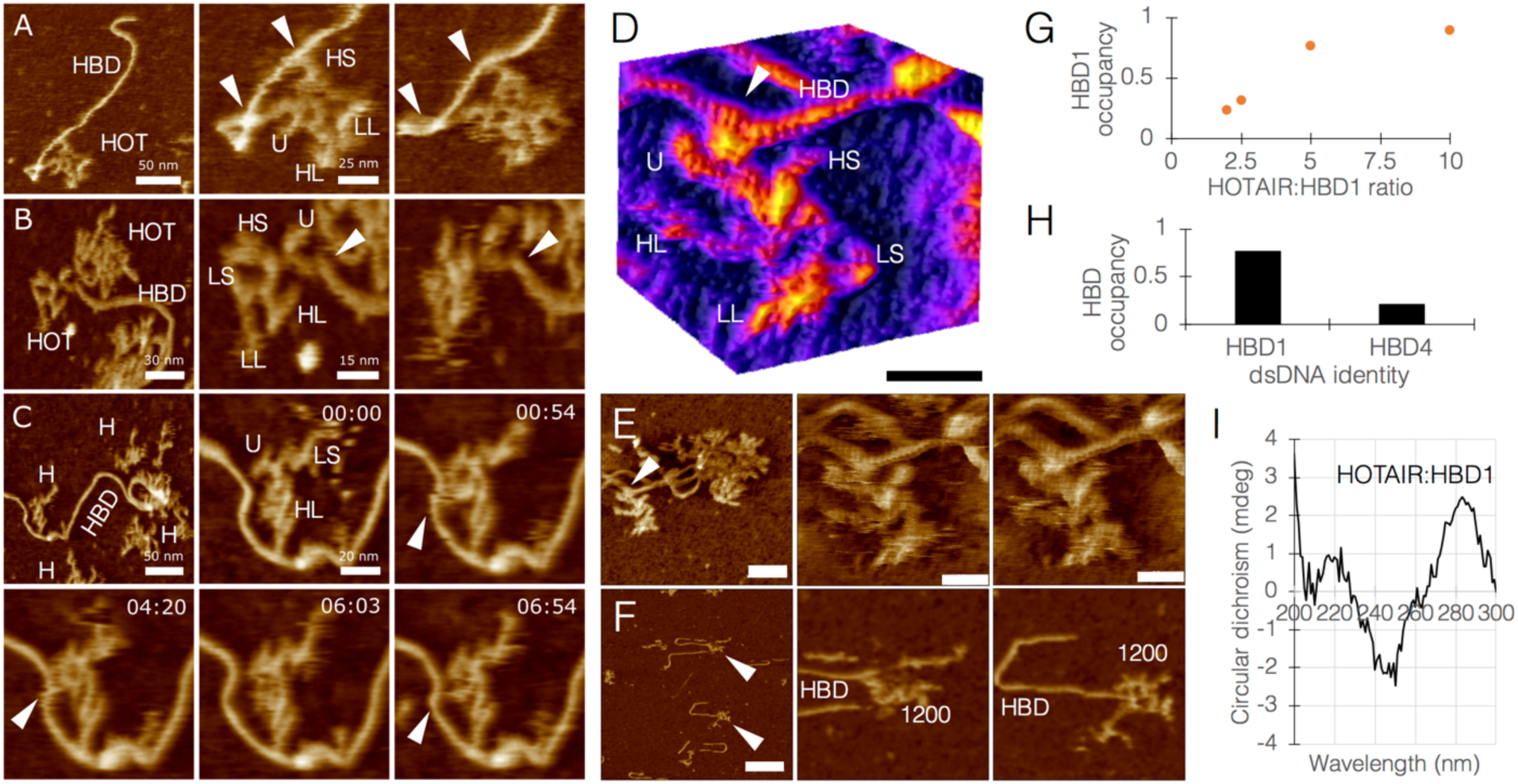
Characterization of the physical interaction between HOTAIR and dsDNA. A-C, AFM images of HOTAIR interacting with HOTAIR-binding DNA 1 (HBD1). White arrowheads point at sites of interaction. (A, B) Center and right panels represent pre-and at-interaction states, recorded with a time difference of 4 min 17 sec, and 17 min 43 sec respectively. C, time series showing a live interaction. HOT, HOTAIR. HBD, HOTAIR-binding DNA 1. Reliably-assigned HOTAIR segments are denoted by their abbreviation. HOT(ag) shows HOTAIR molecules in aggregate. D, 3D surface plot of HOTAIR:HBD1 interaction, white arrowhead points at interaction site. The light/dark stripe pattern on HBD and HOTAIR is the actual major/minor grooves (respectively) that build the double helical structure of dsDNA and dsRNA (bar = 20 nm). Image processing details can be found in **Supplementary note 3.** E, raw specimen used for D, HOTAIR and HBD1 currently at interaction. White arrowhead points to the interaction site (left bar = 35 nm; center and right bars = 15 nm). F, binding of the 1200 nt domain of HOTAIR to HBD1 (bar = 70 nm). G, occupancy of HBD1 by various HOTAIR:HBD1 ratios shows ratio dependence (n from 2 to 10 = 73, 240, 60, and 18). H, occupancy of HBD1 vs. HBD4 by HOTAIR at a 5:1 ratio (n = 60 vs. 66). I, circular dichroism spectrum of HOTAIR:HBD1, showing evidence of a physical triplex between ssRNA and dsDNA (negative peak at 210 nm, shifted positive peak at 280 nm).

Interestingly, HBD1 occupancy by the 1200 nt domain was 18 out of 32 total HBD1 counted, suggesting that an additional domain, such as an H segment as observed here, contributes to the higher binding of the whole molecule.

HOTAIR:DNA interactions have been proposed to be mediated through a triple helix structure^13^. A more recent study^26^ showed by electrophoretic mobility shift assay that HOTAIR segments may form RNA–DNA–DNA triplexes, however in a non-biological system (e.g. boiling to 60 °C and cooling). Hoping to shed light on this mechanism, we initially used the Triplexator^28^ package to perform sequence-based predictions of potential sites for triplex formation between HOTAIR and HBD1. Triplexator retrieved 3 potential triplex scenarios, all within the HBD1-binding, 1200 nt domain of HOTAIR. Out of these, 2 were biologically-probable, i.e. sequences with sufficient guanine residues to support Hoogsteen and reverse-Hoogsteen base pairing, and antiparallel configuration. Examination of the highest-scoring combination of oligonucleotides by circular dichroism (CD) spectroscopy showed evidence of a formed triplex involving ssRNA and dsDNA, namely a negative peak at 210 nm and a shifted positive peak at 280 nm **(Fig. 2I, Supplementary note 9)**.

Our observations of HOTAIR revealed a highly flexible molecule, which exhibited a striking diversity of configurations, albeit converging to the same anatomy. A central question is thus whether the known mechanics of RNA allows for such flexibility. Previous works have reported that the flexibility of dsRNA is lower than that of dsDNA, and measurements by orthogonal techniques have yielded persistence length values around 62 nm^29,30^, which is longer than the observed discrete segments of HOTAIR. However, these estimations may not properly reflect the behavior of biological RNA molecules. Computational predictions, as well as recent experimental works that include biochemical methods and direct imaging^8,20^, show that these molecules are rich in unpaired loops of varying sizes, which can be thought of as mechanical joints that allow for the observed flexibility. In order to quantitate this freedom of motion within our experimental system, we measured the range of movement of a single RNA joint in the AFM. We scanned 12-nt RNA joints connected at the edges of DNA origami rectangles used as AFM imaging guides. Segments connected by these joints were able to pivot up to approximately ±100° in the study conditions **(Supplementary note 10)**. Given the fact that, based on biochemical analysis as well as computational predictions applied locally **(Supplementary note 11)**, HOTAIR is rich in such joints, its actual flexibility is likely significantly larger than that predicted for an idealized dsRNA shaft.

We utilized the fast scanning capability of our AFM system in order to quantitate the dynamics of minimally-constrained HOTAIR molecules under biological conditions. The mica surface was covered with poly-L-ornithine in order to pin down, as quickly as possible, the DNA-RNA complexes that formed during the short incubation time. Once the complex is captured by the mica, DNA is no longer moving, whereas RNA appears as softer, smaller, shows a complex 3D structure with multiple non-binding points allowing it to move more freely and adopt a range of possible configurations. Our initial observations revealed a very diverse range of HOTAIR morphologies **(Fig. 3A)**, and further investigation into the dynamics of these molecules demonstrated that this diversity most likely derives from movement, and not degradation or misfolding **(Movies S8-S10)**. All limbs of HOTAIR are capable of pivoting around joints, extending, or retracting **(Fig. 3B. Movie S8)**, including the U-module, which exhibited pincer-like motion and at least one clear joint in segment UL **(Fig. 3C)**. These movements occupied a radius of up to 20 nm from body **(Fig. 3D-E)**.

**Figure 3.**
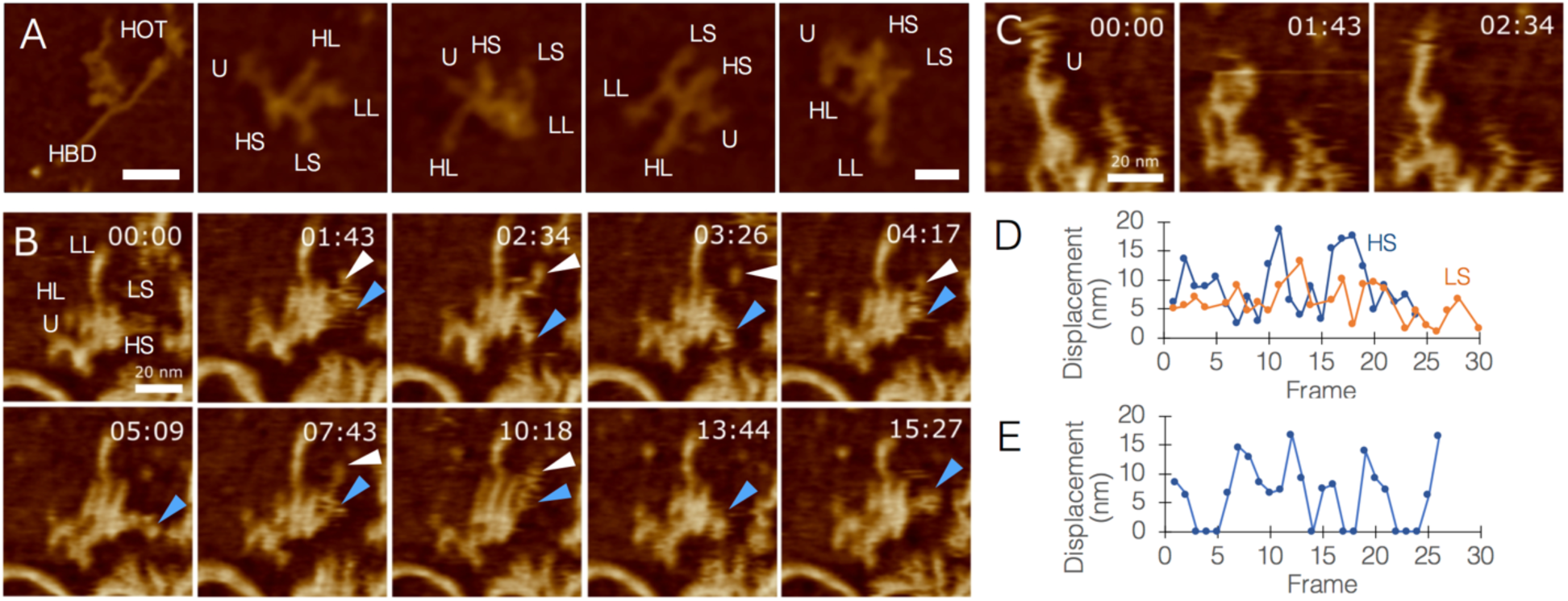
Quantitating the flexibility of HOTAIR. A, a range of HOTAIR morphologies, segment names are abbreviated. Left panel shows HOTAIR interacting with HBD1 through the U-module (left panel bar = 50 nm; other panels bar = 20 nm). B, Timelapse AFM scans of HOTAIR, tracking the movement of segments HS (blue arrowheads) and LS (white arrowheads; bar = 20 nm). C, AFM topography images of HOTAIR focusing on movements of the U-module, exhibiting pincer-like opening/closure and a joint in segment UL (bar = 20 nm). D, Quantitative analysis of the displacement (nm) of HL/HS movements. Numbers were corrected to account for center-mass drift. E, Quantitative analysis of U-module behavior (0 nm represents closed state, other numbers represent varying levels of opening).

Our AFM scans enabled direct observations of structural configurations of DNA and RNA under the study conditions, as well as the identification of differences between their material properties using phase imaging. In this imaging mode, the phase difference between the cantilever and the drive signal is measured, a measurement sensitive to the stiffness or softness of the sample, providing an additional layer of information in samples where height may be equal throughout, but that are made of various materials. Phase imaging of our samples was able to discriminate between DNA and RNA based on the fact that the DNA is more rigid than RNA **(Fig. 4A-B)**. For this reason, we frequently used phase imaging in complex samples to be able to reliably determine molecular identity of sample objects.

**Figure 4.**
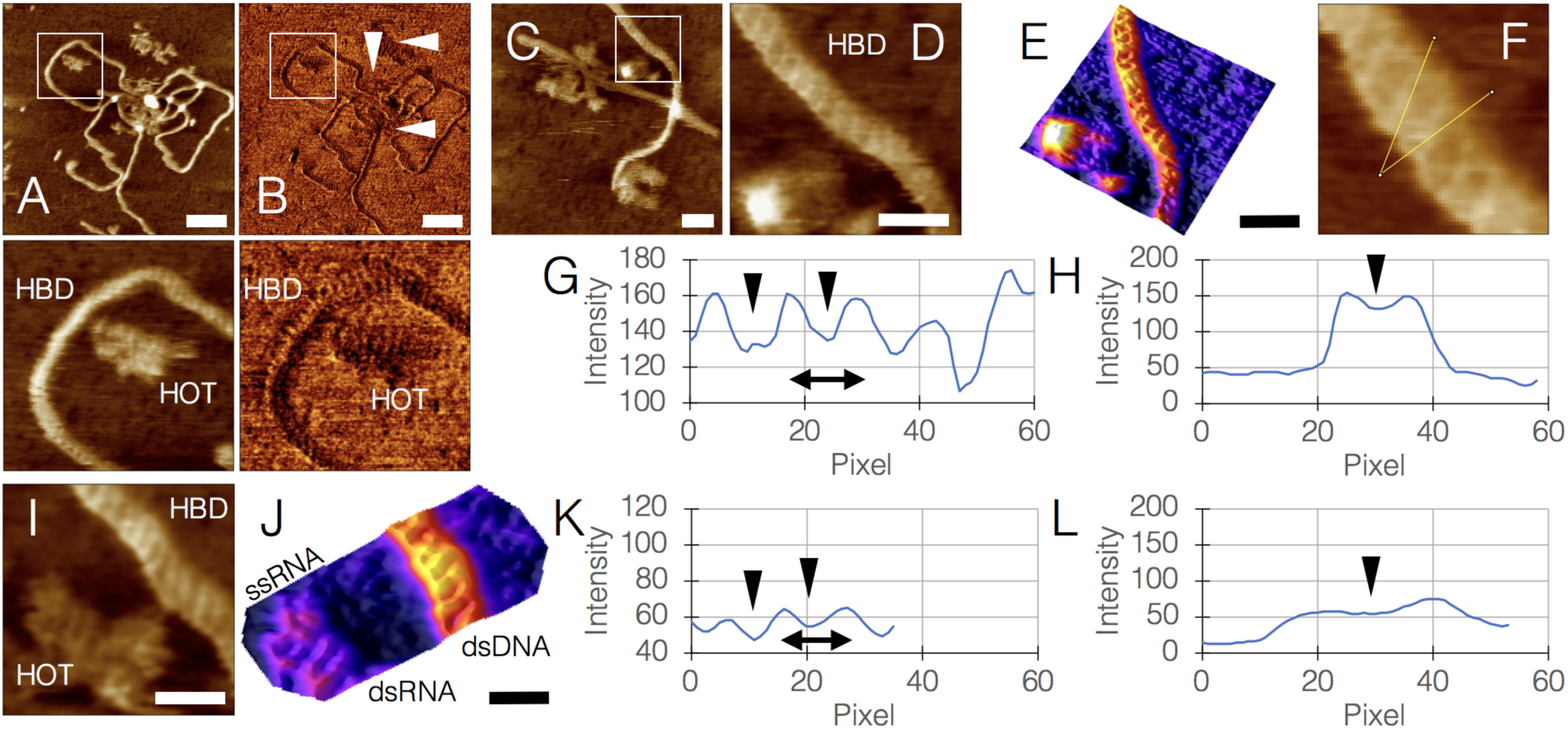
High-resolution imaging of DNA and RNA by AFM. A-B, HOTAIR molecules interacting with HBD1, imaged by height (A) and phase (B), the latter showing differences between HBD1 and HOTAIR molecules based on the higher rigidity of DNA compared with the softer RNA. Bottom panels are magnifications of white squares in A and B (bar = 50 nm) C-D, segment from a HOTAIR:HBD1 sample focusing on the dsDNA structure, D is a magnification of white square in C, showing the right-handed double helix (bar in C = 20 nm. Bar in D = 10 nm). E, 3D surface plot of D (bar = 10 nm). F, helical twist angle measurement, yielding 31.89° relative to helix plane. G, plot profile along the axis showing the major grooves (black arrowheads) and helical pitch (double-sided arrow) of 3.76 nm. H, plot profile orthogonal to the axis. Black arrowhead pointing at oblique segment of a major groove. I, segment of a HOTAIR:HBD1 sample showing dsDNA, dsRNA, and ssRNA together in a single frame (bar = 10 nm). J, 3D surface plot of I, emphasizing the different structures in the sample (bar = 10 nm). K, plot profile along the axis showing the major grooves (black arrowheads) and helical pitch (double-sided arrow) of 2.89 nm. L, plot profile orthogonal to the axis. Black arrowhead pointing at oblique segment of a major groove. Note the lower profile of dsRNA compared with dsDNA (H) due to RNA being softer than DNA.

Multiple scans have repeatedly shown the right-handed, double helical structure of DNA under the study conditions in exquisite detail **(Fig. 4C-E)**. Our measurements yielded a dsDNA helical twist angle of 31.89° relative to helix plane, which is within the observed range^31^ of 27.7°-42.0° **(Fig. 4F)**. Measured mean helical pitch was 3.76 nm, within 3Å of the accepted value of 3.4 nm for B-DNA **(Fig. 4G,H)**. Measurements of dsRNA **(Fig. 4I-J)** yielded a helical twist angle of 18.86° relative to helix plane, within the observed range of 16.1°-44.1°, and a helical pitch of 2.89 nm, with the accepted value being 2.82 nm for A-DNA **(Fig. 4K-L)**. Interestingly, dsRNA exhibited a lower profile than dsDNA due to its relative softness.

In summary, lncRNA molecules are emerging as abundant and important players at multiple levels of regulation over gene expression. Our ability to study structure-function relationships in this new group of molecules would be critical to our understanding of their biology, their roles in health and disease, and the potential ways to correct their malfunction. Our conclusions regarding structure-function relationship of HOTAIR need to be taken carefully as our model does not take into account HOTAIR-protein interactions which may be indispensable to its cellular state and functionality. However, with that said, our observations of HOTAIR produce a first biologically-plausible model of its anatomy, quantitate its plasticity, and confirms it can intrinsically target genomic DNA. Moreover, our findings demonstrates that structural study of lncRNAs can be done using AFM as a tool of choice, owing to its ability to enable direct, high-resolution, and dynamic visualization of nucleic acids in a liquid, cell-like environment, and at physiological temperature. Although it was introduced more than three decades ago^32^, AFM is still not a mainstream technique in molecular, cellular, and structural biology. Our findings make a convincing case in favor of adding this versatile tool to X-ray crystallography, NMR, and cryo-EM, in order to enable new forms of understanding of the behaviors of biological molecules.

## Materials summary

*In-vitro transcription and RNA-seq.* LZRS-HOTAIR and pCDNA3-HOTAIR were a kind gift from Prof. Howard Chang. pJ-HOTAIR was purchased from DNA2.0. IVT templates were either plasmids linearized with EcoRI restriction enzyme (NEB), or PCR amplicons. IVT was carried out using two separate kits (from New England Biolabs and Megascript). IVT was carried out for 3 h at 37 °C and followed by DNA template digestion (using DNase included in the kits). RNA was purified using MegaClear kit. RNA-seq was performed at the Nancy and Stephen Grand Israel National Center for Personalized Medicine (G-INCPM) at the Weizmann Institute of Science. Library preparation was done using in-house protocols.

*AFM and Cryo-EM.* Samples were analysed using a NanoWizard^®^ ULTRA Speed AFM (JPK Instruments, Germany) mounted on an inverted optical microscope (Nikon Eclipse TE2000-U or Zeiss AxioObserver.A1), or equipped with a JPK TopViewOptics™. Samples were imaged in buffer at ambient temperature in amplitude-modulation or phase-modulation AC mode. Fast-scanning high-resonant ultra-short cantilevers (USC-F0.3-k0.3, NanoWorld, Switzerland) with a nominal resonance frequency of 300 kHz in air, spring constant of 0.3 N/m, reflective chromium/gold-coated silicon chip, and high-density carbon tips with a radius of curvature of 10 nm were used. Prior to deposition on substrate, RNA and HBDs molecules were incubated in filtered nuclear-like buffer (NLB; 5 mM NaCl, 140 mM K^+^, 0.5 mM Mg^+2^, 10^-4^ mM Ca^+2^, pH=7.2) for 30 min at 37 °C. For cryo-EM, HOTAIR-bearing grids were plunge-frozen in liquid ethane cooled by liquid nitrogen, using a Leica EM-GP plunger (4 s blotting time, 80% humidity), and imaged at liquid nitrogen temperature on an FEI Tecnai TF20 electron microscope operated at 200 kV with a Gatan side entry 626 cryo-holder. Images were recorded on a K2 Summit direct detector (Gatan) mounted at the end of a GIF Quantum energy filter (Gatan). Images were collected in counting mode, at a calibrated magnification of 16,218 yielding a pixel size of 3.083 Å.

Additional detailed methods can be found in the supplementary notes.

## Acknowledgements

The authors wish to thank M. Gershovits, S. Gilad, Y. Spector and all the staff at the Nancy & Stephen Grand Israel National Center for Personalized Medicine (Weizmann Institute) for discussions and technical assistance, N. Elad (Weizmann Institute) for discussions and technical assistance, F. Buske for kind discussions and technical assistance, S. Dori (Tel-Aviv University) and O. Girshevitz (Bar-Ilan University) for technical assistance, A. Munitz (Tel-Aviv University) for valuable discussions, G. Church (Harvard Medical School) for endless inspiration and valuable discussions, and all the team at Augmanity for valuable assistance and discussions.

## Author contribution statement

RSS spearheaded research. RSS, DS, LA, HH, and JG designed and performed experiments and analyzed data. LA and HJ performed computational modeling. ES and RU analyzed data. IB and AAH oversaw project and designed experiments. IB analyzed data. RSS, DS, and IB wrote the manuscript.

## Competing financial interest

D.M. AND H.H. are employees of JPK, a manufacturer of AFMs. The other authors declare no competing interest in this study.

